# Sex Chromosomes and Gonadal Hormones Contribute to B Cell Antibody Responses in Aging and Alzheimer’s Disease

**DOI:** 10.64898/2026.08.21.746231

**Authors:** Hannah T. Zuppe, Brad T. Casali, Erin G. Reed

## Abstract

It is increasingly appreciated that B cell populations in the brain shift during aging and Alzheimer’s Disease (AD), contributing to neuroinflammation and cognitive decline. Though the underlying mechanisms remain unclear, biological sex is likely a key regulator since differences in B cell antibody responses are one of the most well conserved sex differences in immunology. Sex differences in B cell subtypes and antibody classes are significant because they drive sexually dimorphic immune responses and subsequent disease susceptibility. Despite extensive work evaluating the effects of gonadal hormones on B cells, the effects of aging and AD pathology, and the contributions of sex chromosomes are understudied. Using the Four Core Genotypes (FCG) mouse model to separate chromosomal and gonadal sex, we evaluated B cell subtypes in the brain and bloodstream in both adult and 5xFAD mice via flow cytometry and antibody levels in the cortical brain region of both groups via ELISAs. We found sex differences mediated by both sex chromosomes and the gonadal hormone environment; these differences were primarily involved in antibody class-switching.

## Introduction

While the role of the innate immune system has been extensively studied in Alzheimer’s Disease (AD), the role of the adaptive immune system, primarily mediated by B cells and T cells, remains to be fully understood. Within the adaptive immune system, most of the current research is focused on T cells. However, B cell populations in the brain shift during both aging and neurodegenerative diseases like AD (Kwinta et al. 2026). It is also likely that B cells are contributing to neuroinflammation and cognitive decline in AD, though the exact mechanisms remain unclear.

B cell antibody responses are one of the most well conserved sex differences in immunology (Fink & Klein 2018), indicating biological sex is likely a contributor to B cell-mediated responses in aging and disease. A major component of these sex differences is antibody class-switching, the process through which activated B cells go from producing IgM and IgD to IgA, IgE, or IgG, to effectively combat pathogens. The enzyme Activation-Induced Cytidine Deaminase (AID) breaks at DNA “switch regions” located before the constant regions of the antibody heavy chain. The antigen-binding variable region is then linked to a new constant region for optimal antibody function, and the intervening sequence is excised. Due to AID being activated by estrogen, females typically have higher levels of class-switching (Pauklin et al. 2009), resulting in females having stronger B cell activity, as class-switched antibodies produce more specific immune responses.

While the effects of gonadal hormones on B cells have been extensively studied, the effects aging and AD pathology could have on these differences remains understudied, as is the contribution of sex chromosomes to sex differences in B cell responses generally and AD specifically (Lutshumba et al. 2023). Therefore, we evaluated B cell subtypes and antibody production in the context of aging and AD and whether observed sex differences were primarily due to the sex chromosomes, gonadal hormones, or an interaction of the two. These differences are significant because they drive sexually dimorphic immune responses and subsequent disease susceptibility. In addition, it is potentially therapeutically relevant if an observed sex difference is mediated by sex chromosomes or gonadal hormones.

To determine the source of the observed sex differences, Four Core Genotype (FCG) mice were crossed to 5xFAD mice. In these mice, the *SRY* gene, responsible for testes formation, is deleted from the Y chromosome (Y-) and then inserted onto chromosome 3. This generates XX and XY-testes-bearing and XX and XY-ovary-bearing mice. By comparing mice of the same chromosomal or gonadal sex, it is possible to determine which component is driving an observed sex difference. In the brains of the adult mice, we saw both ovary-bearing and XY-mice had higher percentages of class-switched B cells; this was not seen in the AD mice, likely due to the overall inflammatory milieu driving class-switching. In both adult and AD mice, cortical IgG and IgM levels were higher in both XY- and ovary-bearing mice. This shows that B cell immune responses need the right pairing of chromosomes and gonads, and that the XY-chromosomes-ovaries pairing creates a strongly inflamed B cell phenotype.

## Materials and Methods

### Animal Husbandry

Four Core Genotype (FCG) mice were purchased from The Jackson Laboratory (Strain #010905, originally described in Arnold 2009). These mice carry a mutation in the testes-determining portion of the *Sry* gene on the Y chromosome (Y^−^) that is inserted as a transgene onto chromosome 3, resulting in offspring which may carry the *Sry* transgene, but may possess either testes (XY^−^*Sry* and XX*Sry*) or ovaries (XX and XY^−^). The 5xFAD mouse model (B6.Cg-Tg(APPSwFlLon,PSEN1*M 146L*L286V)6799Vas/Mmjax, RRID: MMRRC_034848JAX, originally described in Oakley et al. 2006) was obtained from the Mutant Mouse Resource and Research Center (MMRRC) at the Jackson Laboratory, an NIH-funded strain repository, and was donated to the MMRRC by Robert Vassar, PhD., Northwestern University. To generate the 5xFAD; FCG mice, XY^−^*Sry* testes-bearing FCG mice were crossed to 5xFAD-transgenic female mice, and offspring were aged to 8 months of age. Mice were ear notched for identification, and tissue was sent to Transnetyx (Cordova, TN) for genotyping using proprietary probes. Animals were housed in microisolator barrier housing with access to food and water ad libitum and on a 12-h light/dark cycle. All animal procedures were approved by the Institutional Animal Care and Use Committee at NEOMED.

### Tissue Collection

Animals were deeply anesthetized with isoflurane using a vaporizer and then euthanized via cervical dislocation.

For brain flow cytometry, the whole brain was manually homogenized on ice in HBSS (Thermo Scientific #14025076) and then held on ice in a heat-activated digestion buffer comprised of Collagenase D (Sigma-Aldrich #11088858001), DNase I (Roche #10104159001), and HBSS until all samples were collected. Samples were incubated at 37°C for thirty minutes and homogenates were stabilized in 2.0 mM EDTA/PBS and filtered through 30 µm filters. Cells were isolated with a 70%-30% Percoll gradient. A single cell suspension was obtained by passing through 30 µm filters a second time.

For blood plasma flow cytometry, whole blood was collected into an EDTA-coated syringe and held on ice until all samples were collected. Blood was incubated in red blood cell (RBC) lysis buffer (BioLegend #420301, diluted 1:10) at room temperature for at least ten minutes in the dark. Blood was spun down in RBC lysis buffer three or four times until the supernatant was clear and plasma was collected. A single cell suspension was obtained by passing through 30 µm filters.

For ELISAs, the brain was bisected and the cortex was dissected on ice from one hemisphere, and samples were snap frozen on dry ice and stored at ™80°C until homogenization. Tissue was homogenized using a rotor–stator homogenizer in 4°C tissue homogenization buffer (1 M Tris, 8.5% sucrose [w/v], 0.5 M EDTA and 0.2 M EGTA in ultrapure water, pH 7.4) supplemented with fresh protease inhibitor cocktail (Sigma #P3480; 1:100 dilution). Each ELISA had its own sample aliquot stored at ™80°C until homogenization to minimize freeze-thaw cycles. Total protein determined via BCA normalized the final ELISA data post-assay.

### Flow cytometry

Single cell suspensions from whole brain homogenates and plasma were stained for cell surface markers. First, dead cells were stained with Zombie NIR (BioLegend #423105) on ice in the dark for ten minutes. Cells were then stained with fluorophore-conjugated antibodies on ice in the dark for thirty minutes. The following antibodies were used: CD45-BV510 (BioLegend #103137), CD11b-BV786 (Thermo Fisher Scientific #417-0112-80), CD19-PE Cy7 (BioLegend #152418), IgM-BV605 (BioLegend #406523), and CD43-BV711 (BD #740668). Cells were fixed in 2% PFA on ice in the dark for fifteen minutes. Stained cells were analyzed on a BD FACS Symphony A1 analyzer. FCS Express 7 Research Edition was used for analysis.

### Antibody Isotype ELISAs

Cortical mouse brain homogenates were thawed on ice the day of the ELISA and vortexed for at least thirty seconds to create an even resuspension. Isotypes were determined by kits for IgA (Abcam ab157717, samples diluted 1:50; ab314603, samples diluted 1:30), IgG (Novus Biologicals NBP3-12521, samples diluted 1:75), IgM (Abcam ab133047, samples diluted 1:50) following the manufacturer’s instructions. All antibody concentrations were normalized to total protein determined by BCA. GraphPad Prism 11 was used for data analysis.

## Statistical analysis

All statistical analysis was carried out with GraphPad Prism 11. Grubbs outlier tests (alpha = 0.01) were run for all experiments and outliers were excluded. 2way ANOVAs (alpha = 0.05) comparing chromosomal sex and gonadal sex effects were run for all experiments. p < is denoted as *, p < 0.01 is denoted as **, p < 0.001 is denoted as ***, p < 0.0001 is denoted as ****. When ANOVA results were statistically significant, multiple comparisons between groups of the same chromosomal or gonadal sex were determined with uncorrected Fisher’s LSD post-hoc tests. GraphPad Prism 11 was used for data analysis. Each data point represented an individual mouse. All data were assumed to be normally distributed.

## Results

B cells cross the blood brain barrier while it is still intact (Marchetti & Engelhardt 2020), but the effect that aging and AD could have on B cell subtypes in the brain remains unclear. B cell-mediated immune responses shift during aging and AD (Kwinta et al. 2026), and analysis of the B cell subtypes could help determine if B cell immune responses are dysregulated. We first looked at lymphoid cells (CD45+ CD11b-) as our parent population for B cells (Fig. 1A). We then looked at the percentage of lymphoid cells that were B (CD19+) cells, and which percentages of these B cells were class-switched (IgM-), the innate-like B1 (CD43+), and the adaptive B2 (CD43-) (Fig. 1B). In the 8-month-old adult mice, around 30% of lymphoid cells were B cells; there was a trend towards testes-bearing mice having a higher percentage of B cells though it was not a significant difference (Fig. 1C). In line with class-switching being activated by estrogen, ovary-bearing mice had more class-switched B cells; interestingly, the more significant difference is XY-mice having a higher percentage of class-switched B cells (Fig. 1D). It seems likely that this chromosomal sex difference is estrogen independent, since it has been found that estrogen only increases class-switching in the presence of two X chromosomes (Peckham et al. 2024). There are two broad categories of B cells: B1 and B2. B1 cells quickly secrete IgM antibodies in response to a wide array of pathogens as a first line of defense (Aziz et al. 2015). B2 cells are the conventional B cells that undergo antibody class switching to produce strong, specific responses to pathogens with help from T cells (Hoffman et al. 2016). Most B cells are the adaptive B2 subtype rather than the innate-like B1 (Fig. 1F, 1G).

**Figure 1.**
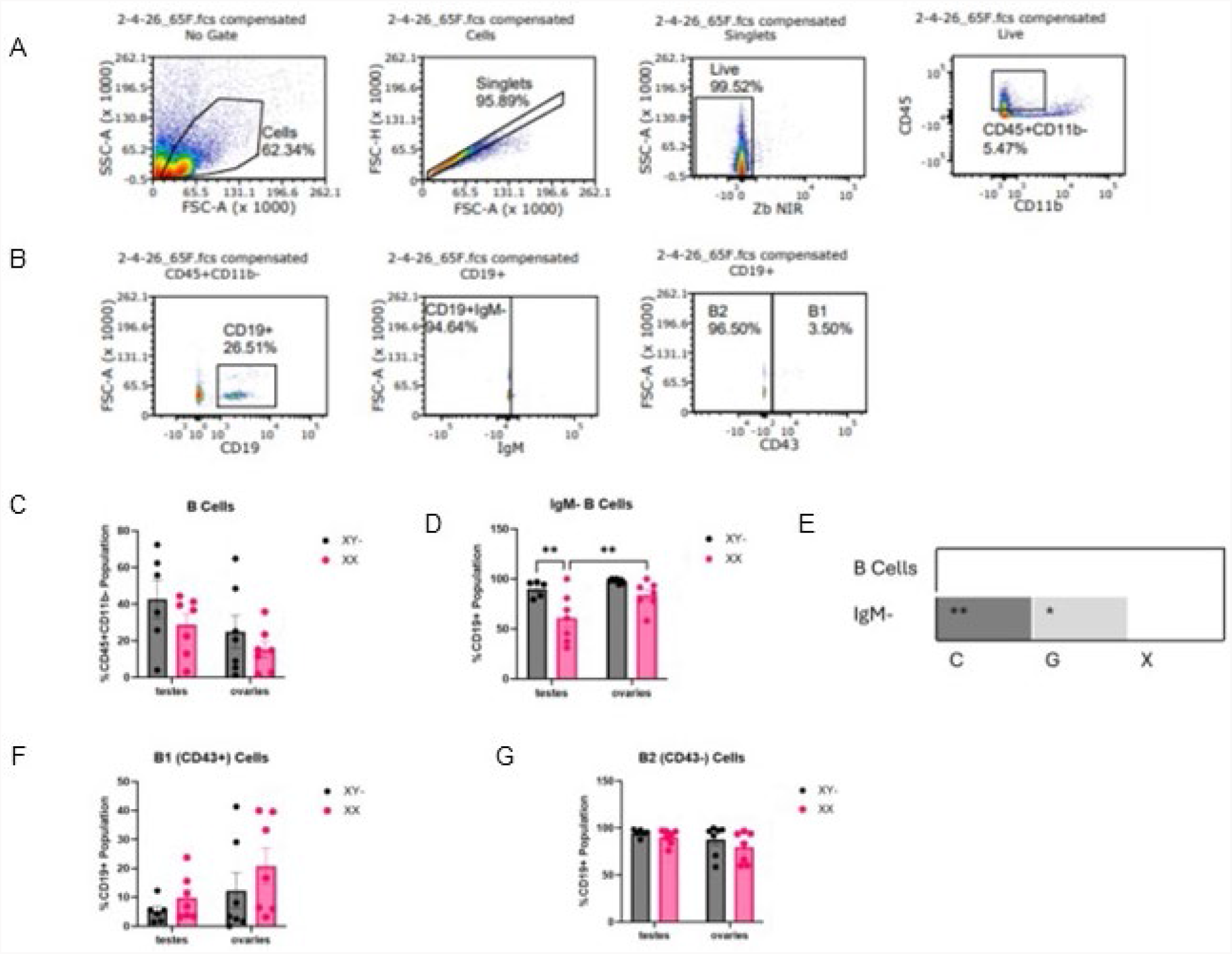
Class-switched B cell levels in 8-month-old wild-type adult mouse brains exhibit distinct chromosomal and gonadal sex differences. Gating strategy for B cells (**A**) and B cell subtypes (**B**) in the brain. (**C**) There are no significant differences in the percentage of B (CD45+CD11b-CD19+) cells from two-way ANOVA, n ≥6/group. (**D**) Percentage of class-switched (IgM-) B cells. Significant pairwise differences from two-way ANOVA are shown, n ≥5/group. (**E**) Summary of two-way ANOVAs determining chromosomal sex (C), gonadal sex (G), and interaction (X) effects. (**F**) There are no significant differences in the percentage of B1 (CD43+) cells from two-way ANOVA, n ≥6/group. (**G**) There are no significant differences in the percentage of B2 (CD43-) cells from two-way ANOVA, n ≥6/group.

Since B cells were present and there were sex differences for class-switching in the adult mouse brains, we repeated flow cytometry in 8-month-old 5xFAD mice to determine if there are AD-specific shifts. Mice were aged to 8 months as they would exhibit robust AD pathology (Westi et al. 2024). There remain a low percentage of B cells comparable to wild-type littermates (Fig. 2A). However, unlike in the wild-type mice, most of these B cells are class-switched (Fig. 2B). This is most likely due to the inflammatory cytokine environment, most notably increased IFN-γ levels, in the AD brains driving antibody class-switching (Paludan 1998, Wang et al. 2016). Most B cells are of the B2, rather than B1, subtype (Fig. 2C, 2D).

**Figure 2.**
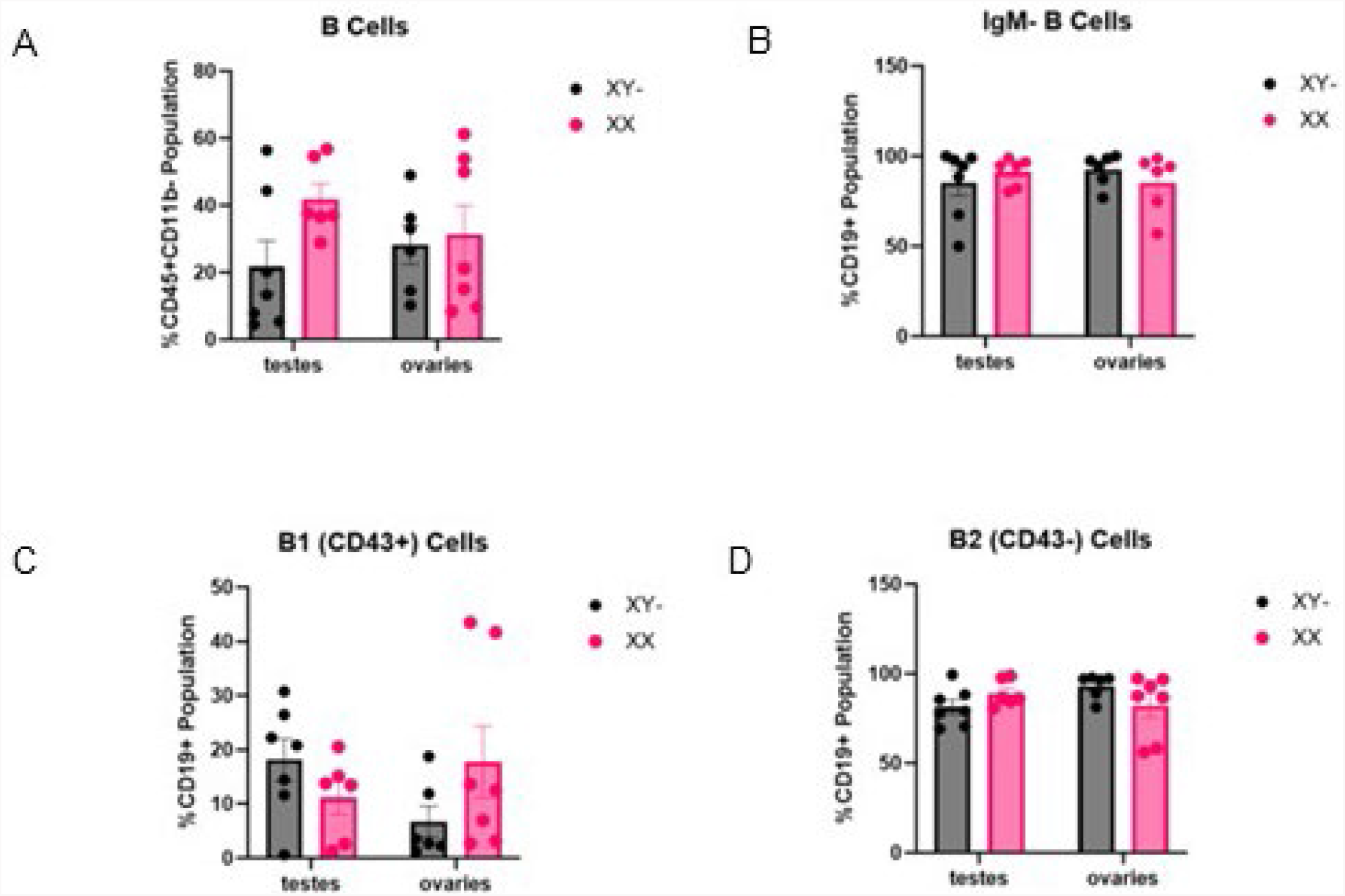
AD brain pathology results in a lack of sex differences in B cell levels and subtypes in 8-month-old 5xFAD mice. (**A**) There are no significant differences in the percentage of B (CD45+CD11b-CD19+) cells from two-way ANOVA, n ≥6/group. (**B**) There are no significant differences in the percentage of class-switched (IgM-) B cells from two-way ANOVA, n ≥6/group. (**C**) There are no significant differences in the percentage of B1 (CD43+) cells from two-way ANOVA, n ≥6/group. (**D**) There are no significant differences in the percentage of B2 (CD43-) cells from two-way ANOVA, n ≥6/group.

B cells secrete antibodies into the cortical brain region with antibodies accumulating as they become less efficient at clearing cellular debris (Frasca, Blomberg 2011). IgA is predominately found in the digestive tract, but it has recently been shown that IgA secreting B cells traffic to the brain during AD (Makhijani et al. 2025). In the cortex of 8-month-old wild-type and 5xFAD mice, IgA concentrations were very low and did not exhibit any sex differences (Fig. 3A, 3E). This is likely due the blood brain barrier still being mostly intact (Yao et al. 2025). IgG accumulates in both aging and AD brains, often binding receptors on microglia to induce an age-associated, pro-inflammatory state (Makhijani et al. 2025). In both wild-type and 5xFAD mice, IgG concentrations are higher in both XY- and ovary-bearing mice, with XY-ovary-bearing mice having the most IgG (Fig. 3B, 3F). This is likely in part due to increased class-switching in the presence of estrogen. While not directly studied, the more inflammatory microglial phenotype in XY-mice could contribute to more IgG (Casali et al. 2025). IgM is a macro-immunoglobulin that compensates for lower antigen binding affinity with multiple binding sites, making IgM particularly adept at binding non-protein antigens (Keyt et al. 2020). Therefore, IgM is important in clearing metabolic byproducts that build up during aging. IgM concentrations are higher in both XY- and ovary-bearing aged mice, with XY-ovary-bearing mice having the most IgM (Fig. 3C). It has recently been found that a gene encoding for IgM, *Ighm*, increases during aging in XY mice and these antibodies bind to carbohydrate waste aggregates in the cortex (Zeng et al. 2026). While it has not been directly studied, it is plausible that IgM is higher in ovary-bearing mice because the age-related shift from carbohydrate to lipid metabolism results in a build-up of carbohydrate aggregates and thus the need for more IgM (Ding et al. 2013). Similar results for IgM are seen in 5xFAD mice (Fig. 3G). Because this metabolic shift is also observed in AD, it may also drive this observation (Ding et al. 2013).

**Figure 3.**
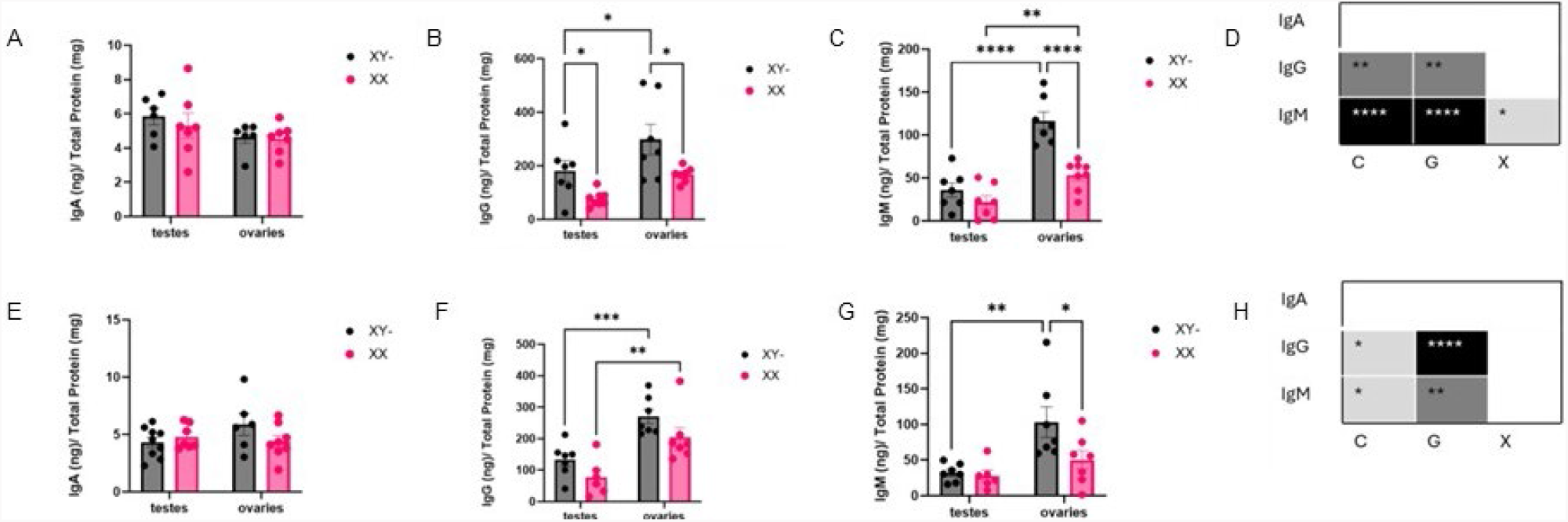
Cortical antibody levels exhibit both chromosomal and gonadal sex differences in 8-month-old wild-type and 5xFAD mice. (**A**) There are no significant differences in IgA concentrations in wild-type mice from two-way ANOVA, n ≥6. (**B**) IgG concentrations from wild-type mice. Significant pairwise differences from two-way ANOVA are shown, n ≥6/group. (**C**) IgM concentrations from wild-type mice. Significant pairwise differences from two-way ANOVA are shown, n ≥6/group. (**D**) Summary of two-way ANOVAs determining chromosomal sex (C), gonadal sex (G), and interaction (X) effects in wild-type mice. (**E**) There are no significant differences in IgA concentrations in 5xFAD mice from two-way ANOVA, n ≥6. (**F**) IgG concentrations from 5xFAD mice. Significant pairwise differences from two-way ANOVA are shown, n ≥6/group. (**G**) IgM concentrations from 5xFAD mice. Significant pairwise from two-way ANOVA are shown, n ≥6/group. (**H**) Summary of two-way ANOVAs determining chromosomal sex (C), gonadal sex (G), and interaction (X) effects in 5xFAD mice.

Before reaching the brain, B cells travel through the blood plasma where their responses will be shaped by both age and sex. At a DNA level, there is a male-specific decrease in B cell specific loci and male-specific shift towards innate immune activity and less adaptive immune activity (Márquez et al. 2020). Additionally, B cell subtypes shift during aging but are context dependent (Frasca 2018). While it has become increasingly apparent that both age and sex modulate peripheral B cell subtypes and overall responses, the interplay between the two remain poorly understood. We repeated the flow cytometry from the brain, sans CD11b as it was used as a marker for microglia (Fig. 4A, 4B). About 40% of immune cells in the blood of 8-month-old wild-type mice are B cells with no observed sex differences (Fig. 4C). Testes-bearing mice have more class-switched B cells in their blood, suggesting that class-switched B cells in ovary-bearing mice have trafficked into tissues rather than staying in circulation (Fig. 4D). Most B cells in circulation are B2 rather than B1 cells (Fig. 4F, 4G).

**Figure 4.**
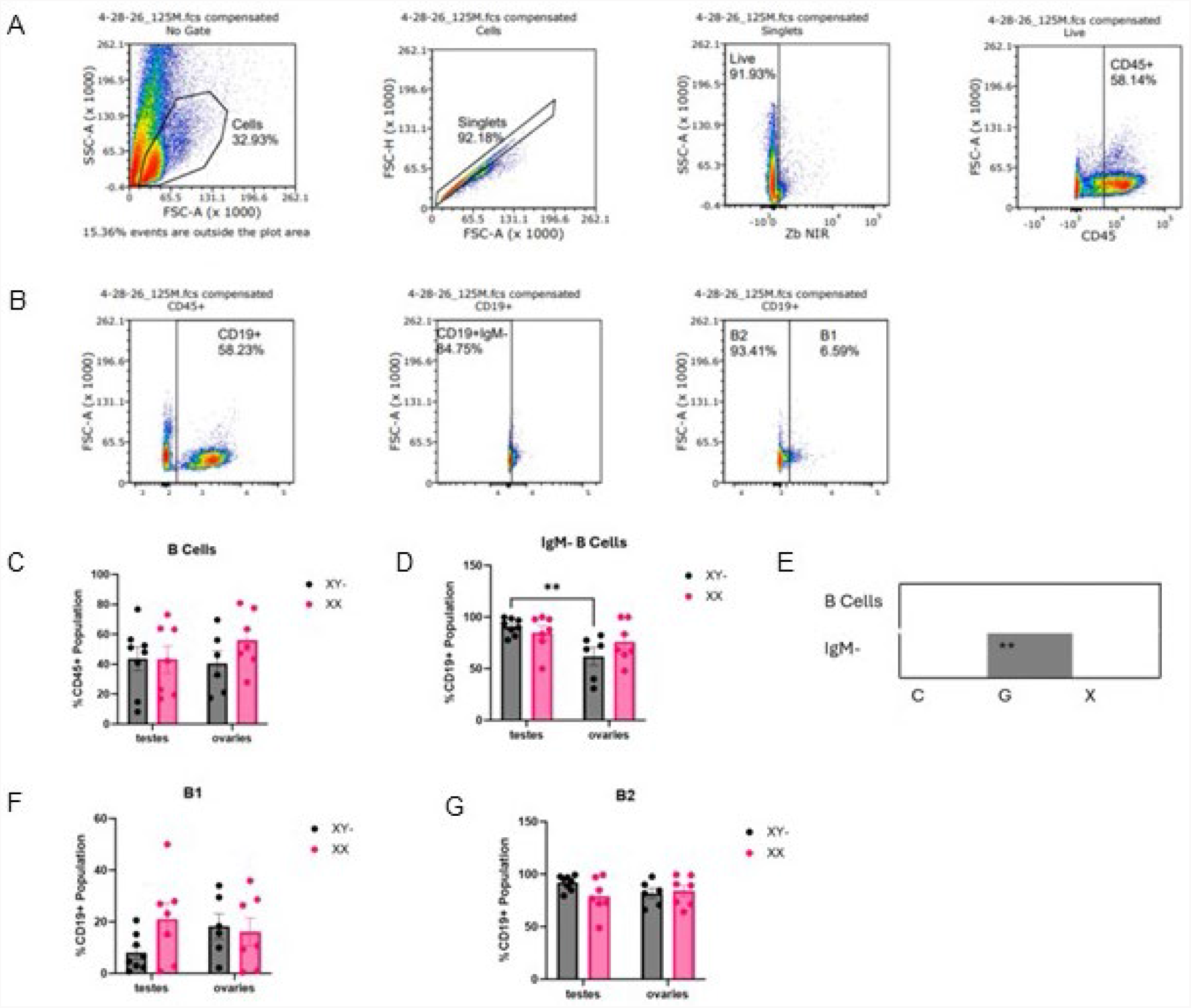
Class-switched B cells in the plasma of 8-month-old wild-type mice exhibit a gonadal sex difference. Gating strategy for B cells (**A**) and B cell subtypes (**B**) in the blood plasma. (**C**) There are no significant differences in the percentage of B (CD45+CD19+) cells from two-way ANOVA, n ≥6/group. (**D**) Percentage of class-switched (IgM-) B cells. Significant pairwise differences from two-way ANOVA are shown, n ≥6/group. (**E**) Summary of two-way ANOVAs determining chromosomal sex (C), gonadal sex (G), and interaction (X) effects. (**F**) There are no significant differences in the percentage of B1 (CD43+) cells from two-way ANOVA, n ≥6/group. (**G**) There are no significant differences in the percentage of B2 (CD43-) cells from two-way ANOVA, n ≥6/group.

Antibodies in the bloodstream of AD patients have been of interest as a diagnostic marker since anti-Amyloid β (Aβ) antibodies, particularly Aβ42, are increased in the blood of AD patients (Söllvander et al. 2015). However, these antibodies can be difficult to accurately measure as they are part of Aβ aggregates making the B cells themselves an appealing target (Söllvander et al. 2015). As circulating B cells in AD remain poorly understood, we repeated the flow cytometry analysis with 8-month-old 5xFAD mice. About 40% of white blood cells in the bloodstream of AD mice are B cells (Fig. 5A). Similar to what was observed in the brain, most of these B cells are class-switched (Fig. 5B). Since class-switching occurs primarily in the spleen, cells travel through the blood to the brain and thus brain and blood results should match. While most B cells are the B2 subtype rather than B1, there is an interesting interaction effect that is only observed in AD. When the chromosomal and gonadal complement does not match (i.e. XX testes-bearing mice and XY-ovary-bearing mice), there is a higher percentage of B1 cells accompanied by a subsequently lower percentage of B2 cells (Fig. 5C, 5D). As B1 cells are a major producer of anti-Aβ IgM, more B1 cells may protect against cognitive decline by producing more anti-Aβ IgM and thus maintaining higher protective antibody levels (Baulch et al. 2020).

**Figure 5.**
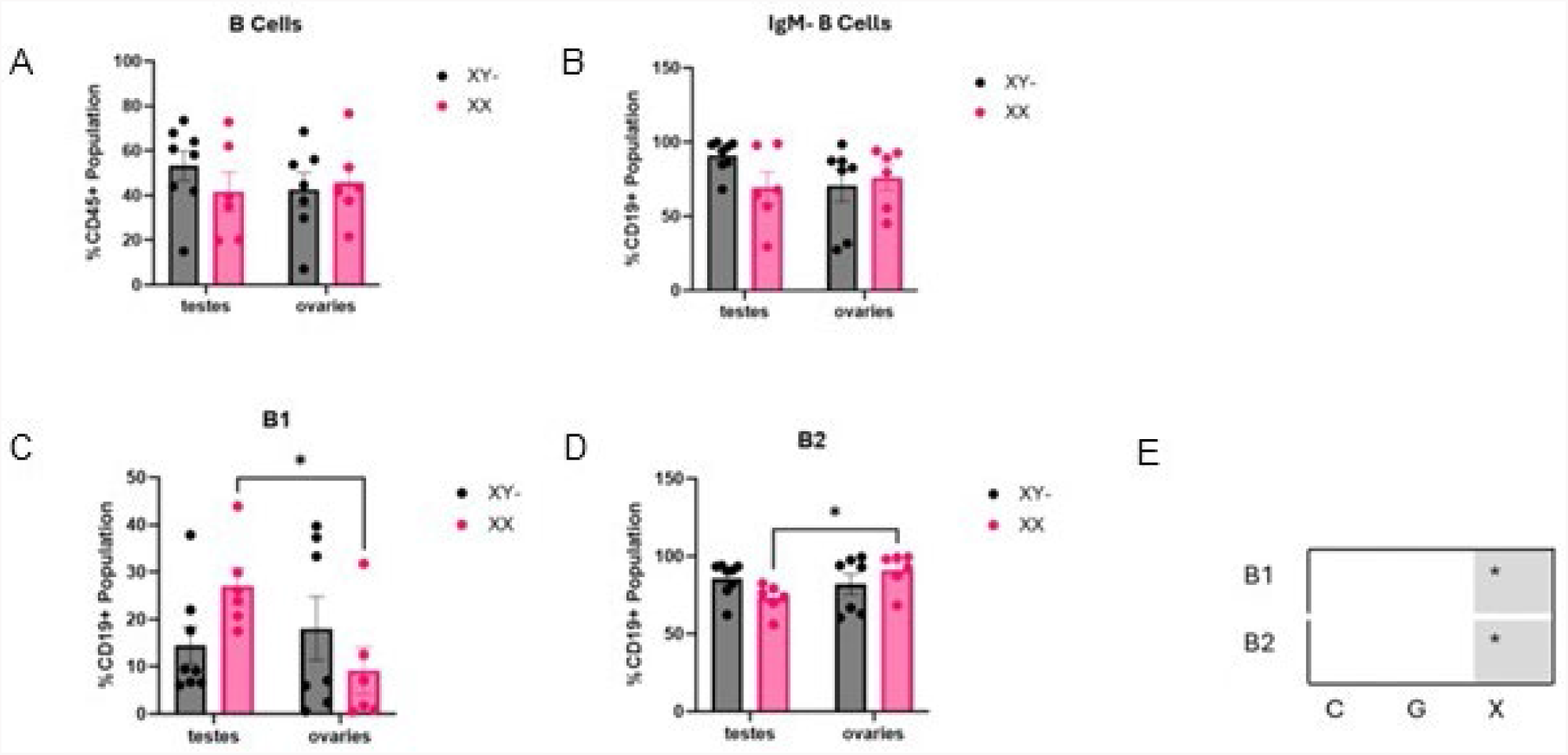
Class-switched B cells in the blood of AD mice exhibit a gonadal sex difference. (**A**) There are no significant differences in the percentage of B (CD45+CD19+) cells from two-way ANOVA, n ≥6/group. (**B**) There are no significant differences in the percentage of class-switched (IgM-) B cells from two-way ANOVA, n ≥6/group. (**C**) Percentage of B cells that are B1 (CD43+). Significant pairwise differences from two-way ANOVA are shown, n ≥6/group. (**D**) Percentage of B cells that are B2 (CD43-). Significant pairwise differences from two-way ANOVA are shown, n ≥6/group. (**E**) Summary of two-way ANOVAs determining chromosomal sex (C), gonadal sex (G), and interaction (X) effects.

## Discussion

In this study, we present an analysis of B cell subtypes in the brain and blood of mice with robust AD pathology and aged mice. In the brains of the aged mice, both ovary-bearing and XY-mice had higher more class-switched B cells. This seems to be independent of the previously known difference due to AID enzyme activation by estrogen since if it was solely mediated by AID activation only XX ovary-bearing mice would have a higher percentage of class-switched B cells (Peckham et al. 2024). It is possible that the XY-mice have a higher percentage of class-switched B cells trafficking to the brain specifically, though the only difference in blood was that testes-bearing mice seemed to have a higher percentage of class-switched B cells staying in circulation. These differences were not seen in the AD mice, likely due to the overall inflammatory milieu, particularly increased peripheral levels of IFN-γ, driving class-switching (Huang 2026).

We also looked at the cortical antibody environment in both AD and aged mice. In both AD and aged mice, IgG levels are higher in both XY- and ovary-bearing mice with XY-ovary-bearing mice having the highest IgG levels, possibly in an inflammatory feedback loop with microglia that exhibit similar sex differences (Casali et al. 2025). Like IgG, IgM levels are higher in both XY- and ovary-bearing mice with XY-ovary-bearing mice having the highest levels, though it seems more likely that IgM levels are being influenced by the brain’s metabolic shifts. It is known that *Ighm*, the gene encoding for the constant region of the IgM heavy chain, increases during aging in XY mice, likely related to the accumulation of carbohydrate aggregates in the cortex (Zeng et al. 2026). Taken together, this suggests that proper B cell function requires matching chromosomes and gonads, while an XY-chromosome and ovary pairing causes severe B cell inflammation.

Limitations of this study were mostly related to not having gonadectomized controls that would ensure that chromosomal differences were not indirectly influenced by gonadal hormones and could have ensured that gonadal hormone effects were not observed gonadectomized controls. However, it is still important to investigate how sex chromosomes and gonadal hormones interact; in addition, these controls were not used because in the context of aggressive AD mouse models, gonadectomies often lead to sexually dimorphic changes in amyloid accumulation and more severe cognitive impairment especially in female mice (Song et al. 2025, Muñoz et al. 2026).

## Ethics approval

All animal procedures were approved by the Institutional Animal Care and Use Committee at NEOMED.

## Competing interests

The authors declare they have no competing interests.

## Funding

This work was supported by the National Institute on Aging of the National Institutes of Health (Award Number R01AG075897 to E.G. Reed), the donors of Alzheimer’s Disease Research, a program of BrightFocus Foundation (A2021036S to E.G. Reed) and institutional funds (to E.G. Reed).

## Notes

### Competing Interest Statement

The authors have declared no competing interest.

